# ProSynTax: A curated protein dataset for taxonomic classification of *Prochlorococcus* and *Synechococcus* in metagenomes

**DOI:** 10.1101/2025.03.20.644373

**Authors:** Allison Coe, James I. Mullet, Nhi N. Vo, Paul M. Berube, Maya I. Anjur-Dietrich, Eli Salcedo, Sierra M. Parker, Konnor VonEmster, Christina Bliem, Aldo A. Arellano, Kurt G. Castro, Jamie W. Becker, Sallie W. Chisholm

## Abstract

*Prochlorococcus* and *Synechococcus* are abundant marine picocyanobacteria that contribute significantly to ocean primary production. Recent genome sequencing efforts, including those presented here, have yielded a large number of high-quality reference genomes, enabling the classification of these picocyanobacteria in marine metagenomic sequence data at high phylogenetic resolution. When combined with environmental data, these classifications can guide cluster/clade/grade assignments and offer insights into niche differentiation within these populations. Here we present ProSynTax, a curated protein sequence dataset and accompanying workflow aimed at enhancing the taxonomic resolution of *Prochlorococcus* and *Synechococcus* classification. ProSynTax includes proteins from 1,260 genomes of *Prochlorococcus* and *Synechococcus*, including single-amplified genomes, high-quality draft genomes, and newly closed genomes. Additionally, ProSynTax incorporates proteins from 41,753 genomes of marine heterotrophic bacteria, archaea, and viruses to assess microbial and viral communities surrounding *Prochlorococcus* and *Synechococcus*. This resource enables accurate classification of picocyanobacterial clusters/clades/grades in metagenomic data – even when present at 0.15% of reads for *Prochlorococcus* or 0.03% of reads for *Synechococcus*.

## BACKGROUND / SUMMARY

*Prochlorococcus* and *Synechococcus* are oxygenic photosynthetic bacteria that are key microbes at the base of the marine microbial food web. Together, they have colonized much of the surface ocean^1^, with *Synechococcus* having a broad geographic range and *Prochlorococcus* dominating the tropical and subtropical open ocean. *Prochlorococcus* and *Synechococcus* are part of a diverse community of marine phytoplankton, which have a key role in regulating the transformation of energy and matter in the oceans. Studies of marine ecosystems have generated large metagenomic datasets^2–5^, which are a rich resource for investigating the distributions of microorganisms in the marine environment^6^. Many classification tools have been developed to process growing volumes of sequence data. Some use broad reference databases (e.g., NCBI and GTDB), while others are specialized to specific groups of microbes^7,8^. Accurate species identification is challenging due to the high diversity of natural samples and the limited availability of taxonomically classified genomes, particularly for distinguishing between very closely related organisms.

Determining the distributions of the picocyanobacteria *Prochlorococcus* and *Synechococcus* is of particular importance for understanding marine ecosystems. These organisms are exceptionally diverse^9–13^, thus making the accurate differentiation of clusters/clades/grades within each genus difficult. As a result, *Prochlorococcus and Synechococcus* populations are often classified as a single strain^14^, obscuring the finely-tuned niche partitioning that is well documented in these groups^15–20^. Over the past decade, a substantial amount of high-quality genomic reference data has become available to address these gaps. These data enable the linkage between the functional and taxonomic structure of *Prochlorococcus* and *Synechococcus* populations and the features of the environment in which they are embedded. They also include genomes from uncultivated single cells, dwarfing the number of available reference genomes from cultured cells by an order of magnitude^21–23^. Single-cell genomes have substantially expanded our understanding of picocyanobacterial diversity while mitigating biases inherent in cultivation-dependent methods, which tend to select for strains adaptable to existing isolation protocols.

Here, we leveraged these extensive datasets to present a comprehensive protein dataset – ProSynTax – that has been manually curated to enable finely resolved taxonomic classification of *Prochlorococcus* and *Synechococcus*, along with the microbial and viral communities associated with these picocyanobacteria in short-read metagenomic datasets. The curated taxonomy is based on the core protein similarity of individual genomes as well as decades of research on niche partitioning and the diverse ecology and physiology of different phylogenetic branches of *Synechococcus* and *Prochlorococcus*^12,13,15,17^. While we have chosen to retain this traditional clade/ecotype-based framework for its ecological relevance, the workflow is taxonomy-flexible and allows users to apply alternative classification systems, such as the genome-based taxonomies proposed by Tschoeke et al.^24^ and Salazar et al.^25^, if desired.

Accompanying this protein dataset is a new workflow that links high-throughput quality control of short-read sequences and taxonomic classification using dedicated classifier algorithms^26,27^. This dataset leverages protein reference sequences, demonstrated to perform comparably to nucleotide references for metagenomic classification^28^, and aligns well with our normalization approach, which utilizes single copy core genes. Custom scripts are included for reporting both the absolute and normalized read abundances (the latter referred to as genome equivalents) of *Prochlorococcus* and *Synechococcus* at user-defined taxonomic levels in metagenomic samples, ranging from broad groupings of high-light and low-light adapted *Prochlorococcus* to more fine-grained distinctions between individual clades/grades. The workflow can accurately identify the major clusters/clades/grades/ecotypes of these picocyanobacteria in short-read sequence libraries consisting of a minimum of 1 million reads in which *Prochlorococcus* represents at least 0.80% of reads, and *Synechococcus* at least 0.09% of reads. ProSynTax incorporates protein sequences from 1,260 genomes of *Prochlorococcus* and *Synechococcus*, encompassing single-amplified genomes, high-quality draft genomes, and newly closed genomes. Among its contents are newly closed circular genomes from 39 *Prochlorococcus*, 12 *Synechococcus*, and 12 marine heterotrophic bacterial strains that were co-isolated from *Prochlorococcus* cultures, including 29 previously partially assembled and 10 unpublished isolate *Prochlorococcus* genomes. Furthermore, ProSynTax includes proteins from 41,753 genomes of marine heterotrophic bacteria, archaea, and viruses to analyze the microbial and viral communities associated with *Prochlorococcus* and *Synechococcus*. The closed genomes reported here will be particularly valuable for studies of microbial evolution that rely on fully assembled genomic islands and regions of the genome that are subject to remodeling.

## METHODS

### Enrichment and isolation of picocyanobacteria

Metadata for culture isolates included in this dataset, including both newly closed genomes and previously published partial genomes that were missing detailed isolation information, are defined here. Specifically, the strains were isolated from seawater collected from 3 cruises in the North Pacific Subtropical Gyre and 1 cruise in the South Pacific Subtropical Gyre. Ten low-light (LL) adapted *Prochlorococcus* strains representing the LLI and LLIV clades and 4 heterotrophic bacteria representing classes Gammaproteobacteria and Alphaproteobacteria were isolated and sequenced (Fig. 1, Table 1). These 4 marine heterotrophic bacteria were co-isolated from *Prochlorococcus* cultures. *Synechococcus* strain Cu2B8 was also isolated and sequenced; however, records of its isolation location and depth are not available (Table. 1).

**Table 1.**
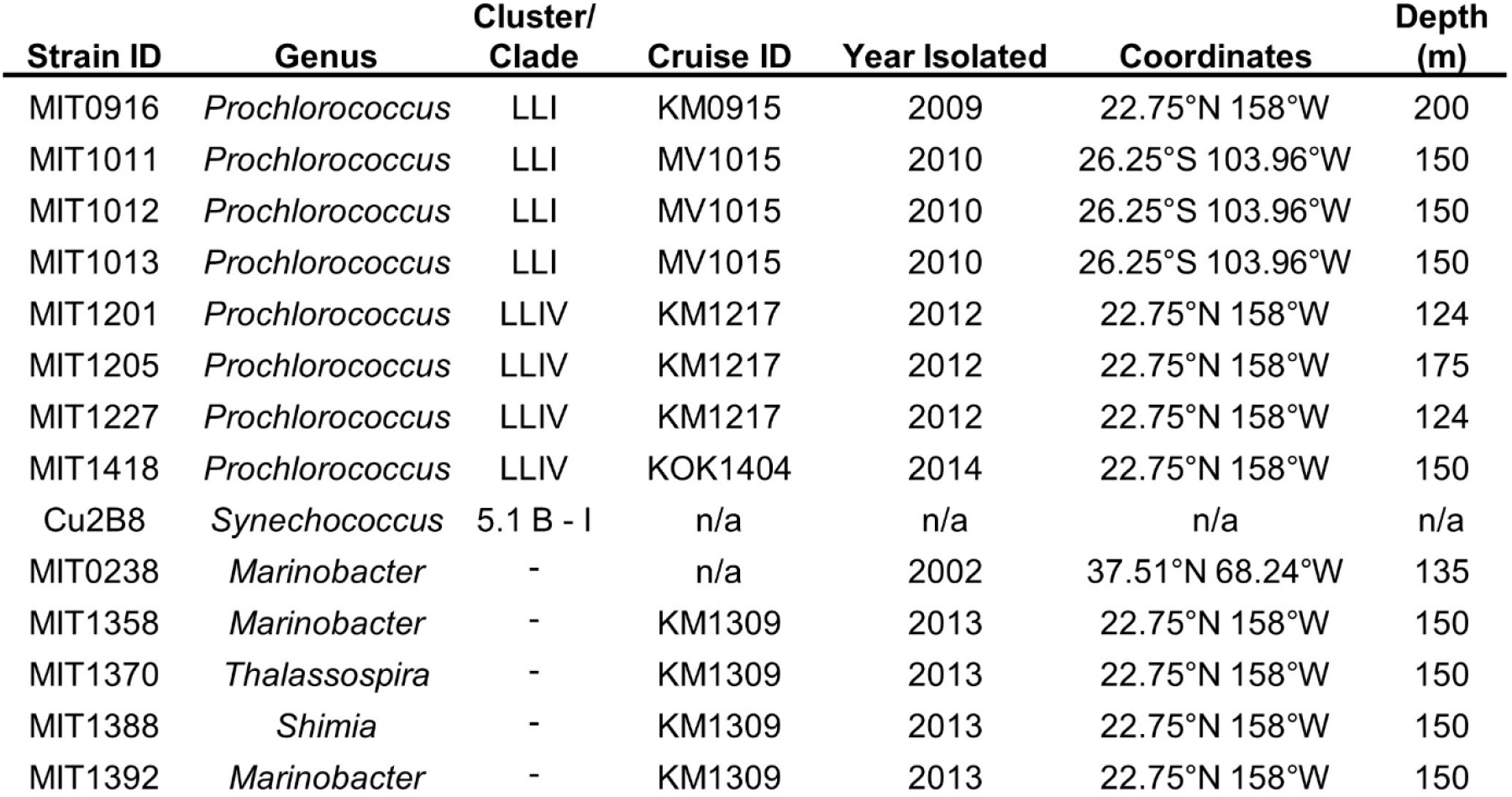
Culture isolates of *Prochlorococcus, Synechococcus*, and marine heterotrophic bacteria. Clade and cluster designations for *Prochlorococcus* and *Synechococcus* strains are provided, along with isolation location, depth, year, and cruise identifier.

**Figure 1.**
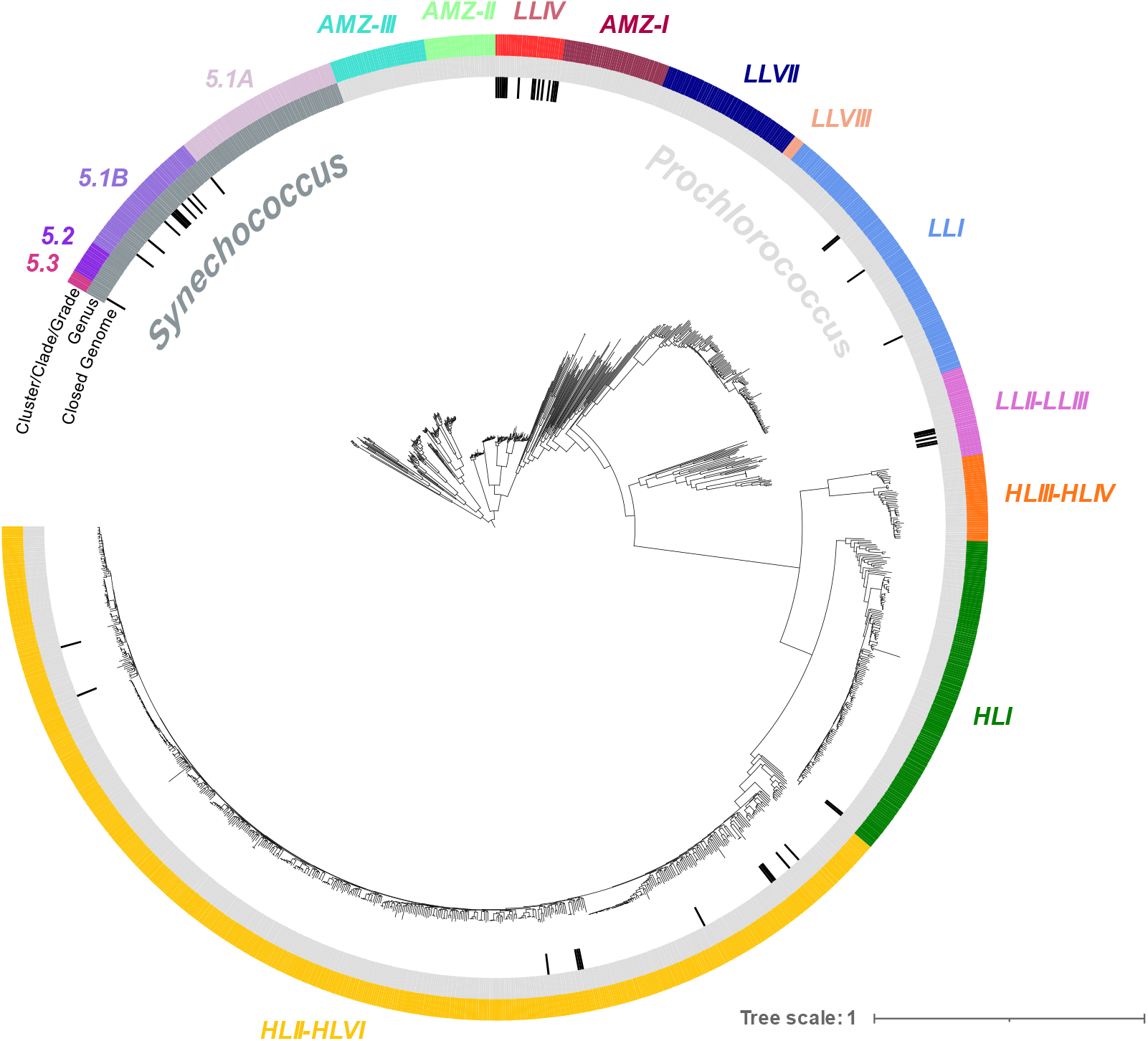
Phylogenetic tree of *Prochlorococcus* and *Synechococcus* included in ProSynTax. The phylogeny includes 1,106 *Prochlorococcus* genomes and 154 *Synechococcus* genomes and is based on a concatenated protein alignment of the 424 single-copy core genes (see Materials and Methods). The tree is arbitrarily rooted at the *Synechococcus* cluster. The outer layer (colors) depicts the major cluster, clade, or grade defined in ProSynTax for each genome. The middle layer (shades of gray) contains the associated genus for each genome, and the inner layer (black bars) highlights closed genomes that are presented for the first time in this work.

All *Prochlorococcus* strains were isolated from raw seawater, except MIT1227 and MIT1418 which were isolated from the filtrate of seawater put through a 1 or 1.2 µm filter. Different media formulations were used, with specific nutrient amendments aimed at selecting for various clades of *Prochlorococcus*, as described hereafter. Isolate MIT0916 was amended with Pro2 nutrients^29^, replacing the nitrogen source with 50 µM nitrate. Isolates MIT1011, MIT1012, and MIT1013 were isolated on media containing 20 µM ammonium chloride, 1 µM sodium phosphate, 0.1x Pro99 trace metal mix^29^, and 1 mM sodium bicarbonate. Isolates MIT1201 and MIT1205 were isolated on media containing 16 µM ammonium chloride, 1 µM sodium phosphate, and 0.1x Pro99 trace metal mix. MIT1227 was isolated on media containing 32 µM ammonium chloride, 2 µM sodium phosphate, and 0.1x Pro99 trace metal mix. Finally, MIT1418 was isolated on media containing 0.1x Pro2 nutrient and metal concentrations. The chemicals used in media preparation were of the highest purity (i.e. Sigma BioUltra) to prevent contaminants that could affect culture growth.

Heterotrophic bacteria MIT1350, MIT1358, MIT1370, MIT1388, and MIT1392 were isolated on 0.3% agar pour plates consisting of Pro99 medium amended with 0.05% pyruvate and 3.75mM TAPS buffer. Colonies were picked and subcultured three times before being transferred into liquid Pro99 medium. MIT1350 was sequenced before freezing, while the remaining strains were directly frozen. Upon thawing MIT1358, MIT1370, MIT1388, and MIT1392 for this study, the cells were repurified using the same methods, with the exception that colonies were transferred into liquid ProMM medium^30^ before pelleting. Purity was then assessed through genomic sequencing to verify the absence of contaminants. Heterotrophic bacteria strain MIT0238 was isolated from xenic *Prochlorococcus* MIT9313; however, the records detailing this isolation are not available.

### Culture conditions and biomass preparation

Cultures were grown in 200 mL of Pro99 medium^29^ at 24°C using continuous light or a 13:11 light-dark cycle at light intensities ranging from 10 to 50 µmol photons m^-2^ s^-1^. Cells were harvested for sequencing during late exponential growth phase, with 0.003% Pluronic F-68 Polyol (MP Biomedicals, Cat# 2750049) added to promote cell aggregation and improve biomass yields during centrifugation. Cells were centrifuged at 7,197 *x g* for 15-20 min at room temperature. After centrifugation, the supernatant was decanted, and the resulting pellets were flash frozen in liquid nitrogen and stored at -80°C.

### Extraction and sequencing

Genomic DNA was extracted from the frozen cell pellets using a modified version of a phenol:chloroform based extraction method^31^ in order to obtain high molecular weight DNA suitable for sequencing on Pacific Biosciences (PacBio) platforms. Modifications included minimizing agitation to prevent DNA shear (e.g., gently shaking the sample tube by hand instead of vortexing), adding 1µl of 50 mg/mL lysozyme (cat# 90082, Thermo Scientific) to the cell pellet during the digestion step, performing a second chloroform/isoamyl alcohol step to remove residual phenol, and increasing the salt concentration by the addition of 0.1 volume of 3M sodium acetate or 3M ammonium acetate before precipitation with 0.6-0.8 volumes of isopropanol. The tube was then gently inverted multiple times and incubated at room temperature for 30 min. DNA was recovered by centrifugation at 10,000 x *g* for 3 min at 4°C. The supernatant was then carefully removed by pipetting. The pellet was then washed twice with 75% ethanol before allowing the pellet to dry at room temperature for approximately 10 min inside a biological safety hood to prevent contamination. It was then dissolved in 1x TE buffer (pH 8) and stored at -80°C.

Genomic DNA samples were diluted and fragmented to 10-12Kb using a gTube (cat# 520079, Covaris), followed by a 0.45X SPRIselect (cat# B23317, Beckman Coulter Life Sciences) bead cleanup. The samples were then made into indexed SMRTBell libraries following guidelines for the template prep kit v 1.0, 2.0, or 3.0 (cat# 100-259-100, 100-938-900, 102-141-700, respectively, Pacific Biosciences). These libraries were assembled into a single pool, and cleaned using MinElute reaction cleanup kit (cat# 28204, Qiagen), followed by additional cleaning using 0.4X SPRIselect beads. The libraries were then bound using Sequel Binding Kit v 2.1, Sequel II Binding Kit v 2.1, and Sequel II binding Kit v 3.2 (cat#100-369-800, 101-820-500, 102-194-100, respectively, Pacific Biosciences) and sequenced on PacBio Sequel I, Sequel II, or Sequel IIe flowcells at either the MIT BioMicro Center or Harvard Bauer core facilities.

### Genome assembly

PacBio sequence data were demultiplexed by barcode using PacBio SMRTtools, generating both circular consensus sequencing reads and subreads. To maximize the likelihood of closing genomes, reads were assembled using multiple tools, including Flye v2.9.5, Canu v2.3, and SPAdes v4.1.0^32–34^. Contigs were classified using MMseqs2 v17.b804f against the GTDB v207 genome database^35,36^. The resulting genomes were submitted to NCBI and IMG, with accessions provided in ProSynTax_genomes.csv^37^. The GitHub repository, which includes documentation and code for genome assembly, is provided in the Code Availability section.

In addition to closing genomes of cultured strains, six heterotrophic bacterial genomes were successfully assembled from sequencing xenic cultures of *Prochlorococcus* and *Synechococcus*. However, since pure axenic isolates for these heterotroph strains are unavailable, we implemented a new nomenclature by appending an additional two-digit identifier (“01”) to the picocyanobacterial strain name from which they were derived. For example, *Thalassospira* MIT121401 was obtained from sequencing a xenic culture of *Prochlorococcus* MIT1214.

### Dataset construction

The dataset consists of single-amplified genomes, finished genomes of culture isolates, and high-quality draft genomes of cultured isolates. These genomes were obtained from several sources, including IMG-ProPortal^21^, GORG-Tropics^38^, NCBI RefSeq (July 2025), Cyanorak^39^, and this study^37^. No metagenome-assembled genomes were included in the dataset to ensure only high-quality reference genomes were present. Additionally, we filtered the NCBI RefSeq database to include all viral genomes and only bacterial and archaeal reference genomes, which typically includes a single reference genome per species. For any genomes without annotated protein sequences, prodigal v2.6.3^40^ was used to identify open reading frames. Stop codons were removed from these predicted protein sequences to remain consistent across all genomes. All protein sequences were concatenated, and Kaiju v1.10.1 was used to construct the finalized dataset (kaiju-mkbwt and kaiju-mkfmi with default parameters)^26^.

### Picocyanobacteria taxonomy

The taxonomy of *Prochlorococcus* and *Synechococcus* was manually curated based on a concatenated multiple-sequence alignment of proteins encoded by any single-copy core genes annotated in each genome. Single-copy core genes were defined as those present in exactly one copy (i.e., genes with multiple copies within any genome were removed) in 100% of 92 genomes derived from cultures in the CyCOG v6.0 database^21^ and estimated to be >99% complete using checkM^41^. All proteins encoded by *Prochlorococcus* and *Synechococcus* genomes were aligned against proteins in the CyCOG v6.0 database (blastp v2.16.0 -evalue 0.001), selecting the best hit for each gene. Following the annotation of all genomes using the CyCOG v6.0 database, protein sequences for the 424 single-copy core genes were aligned using ClustalOmega v1.2.4^42^, concatenated with MEGA v11.0.13^43^, and used to construct a phylogenetic tree (Fig. 1) with FastTreeMP v2.1.11 (-lg -boot 100)^44^. Genomes missing more than 95% data among positions with <50% gaps or genomes with less than 5 single-copy core genes were removed from the dataset and phylogenetic tree. The classified genomes were then analyzed using CheckM v1.2.3^41^ to determine the completeness and contamination for each genome.

## DATA RECORDS

ProSynTax data files are accessible online through a Zenodo repository^37^, which includes the following files:

### ProSynTax_genomes.csv

Table of genomes included in the ProSynTax dataset and their associated metadata. Data fields are as follows:

organism: The name of the organism recorded in NCBI when available. For genomes/organisms obtained from sources other than NCBI, the organism’s name is provided in NCBI format

genome_short_name: The genome name used in the ProSynTax dataset

domain: Bacteria, Archaea, Eukarya, or Virus

genus: The genus of the organism in NCBI

clade: The major cluster/clade/grade of *Prochlorococcus* or *Synechococcus* based on phylogenetic reconstruction using a concatenated alignment of proteins encoded by single-copy core genes

NCBI_BioProject: The NCBI BioProject accession number associated with the organism, when available

NCBI_BioSample: The NCBI BioSample accession number associated with the organism, when available

NCBI_GenBank: The NCBI GenBank accession number associated with the genome sequence data, when available

IMG_Genome_ID: The IMG Genome ID accession number, also known as the IMG Taxon ID, corresponds to the genome or organism in the Joint Genome Institute’s (JGI) Integrated Microbial Genomes (IMG) repository, when available

### ProSynTax_names.dmp

Names taxonomy file for use with the ProSynTax dataset

### ProSynTax_nodes.dmp

Nodes taxonomy file for use with the ProSynTax dataset

### ProSynTax_v1.1.fmi.bz2

Index file containing contents of ProSynTax_v1.faa for use with ProSynTax dataset

### CyCOG6.dmnd

Database containing orthologous groups of proteins used in the cluster/clade/grade normalization step

### ProSynTax_v1.1.faa.bz2

File containing protein sequences used by Kaiju for classification of reads. Each protein sequence contains a header starting with “>“

### average_cycog_length.csv

Comma separated file containing the average length for each protein sequence used in the normalization step. Data fields are as follows: cycog:

name for single-copy core gene

mean_AA_length: the average length of amino acids in the protein sequence of the gene

### ProSynTax-workflow_benchmarking_genomes.tsv

This tab-delimited file contains a list of subsetted genomes used in each benchmarking experiment done in the Technical Validation section. Data fields are as follows:

Experiment Name: name of benchmarking experiment conducted

Subset ID: unique ID from the random genome subsetting

Genome Name: name of genome used in benchmarking experiment

### ProSynTax-workflow_benchmarking_composition.tsv

This tab-delimited file contains the taxon composition of all samples used in each benchmarking experiment done in the Technical Validation section. Data fields are as follows:

Experiment Name: name of benchmarking experiment conducted

Sample Name: unique sample name

Percent *Prochlorococcus*: percent of reads in simulated sample originating from *Prochlorococcus* genomes

Percent *Synechococcus*: percent of reads in simulated sample originating from *Synechococcus* genomes

Percent Heterotroph: percent of reads in simulated sample originating from marine heterotrophic bacterial genomes

Notes: additional information about the simulated sample

### ProSynTax_v1.1_without_refseq.faa.bz2

File containing protein sequences found in ProSynTax with NCBI RefSeq genomes removed. Each protein sequence contains a header starting with “>“.

## TECHNICAL VALIDATION

### Generation of mock metagenome data

To assess the accuracy of cluster/clade/grade-level classification of metagenomic reads using ProSynTax, we generated mock metagenome sequence datasets with known genome inputs. The mock dataset was generated using a custom script^37^ to randomly select 20% of genomes from each *Prochlorococcus* and *Synechococcus* cluster/clade/grade and 20% of marine heterotrophs from the reference genome dataset for read simulation. These genomes were used as input for mason2 v2.0.0-beta1^45^ to create simulated mock metagenomes with 1 million paired-end 150 nt sequences. The .faa file used for classification was modified to exclude sequences associated with the genomes used in the mock metagenomes. The .fmi file was then rebuilt using the subsetted .faa file using the Kaiju functionalities kaiju-mkbwt and kaiju-mkfmi^26^. This ensures that genomes used to create mock metagenomes were excluded from the set used for classification. Genomes were randomly subsetted 10 times for each technical validation experiment that consisted of a paired mock metagenome and a subsetted taxonomic dataset^37^.

### Assessment of misclassification rates

We evaluated the accuracy of the read classification workflow using the simulated dataset derived from a subset of the reference dataset (described above). Reads for each group were simulated separately, processed through the classification workflow, and evaluated based on the proportion classified as true positives or misclassified as false positives. The average results across 10 subsets are presented in Table 2.

**Table 2.** Misclassification rates of simulated metagenomic reads for *Prochlorococcus, Synechococcus*, and marine heterotrophic bacteria. The data represents the average percent of reads and corresponding standard deviation, derived from 10 randomly sampled genome subsets.

|  |  | Read Group |  |  |
| --- | --- | --- | --- | --- |
|  |  | *Prochlorococcus* | *Synechococcus* | Heterotroph |
| Classified As | *Prochlorococcus* | 81.256 ± 0.451 | 1.507 ± 0.172 | 0.007 ± 0.001 |
|  | *Synechococcus* | 0.777 ± 0.253 | 68.739 ± 2.662 | 0.001 ± 0.001 |
|  | Heterotroph | 2.993 ± 0.127 | 10.077 ± 1.786 | 86.693 ± 0.045 |
|  | Unclassified | 14.974 ± 0.379 | 19.677 ± 1.589 | 13.298 ± 0.046 |

### Limit of Detection

To determine the minimum detectable abundance of *Prochlorococcus* in a sample, we simulated datasets with varying proportions of *Prochlorococcus* and heterotrophs and processed them through the classification workflow. The simulated heterotroph proportions are available on Zenodo (ProSynTax-workflow_benchmarking_composition.tsv)^37^. The *Prochlorococcus* false positive rate was calculated by dividing the number of heterotroph reads misclassified as *Prochlorococcus* by the total reads classified as *Prochlorococcus*. To maintain a misclassification rate below 10%, *Prochlorococcus* abundance must exceed 0.08% of all classified reads (Fig. 2 A). Using the same approach, *Synechococcus* requires an abundance greater than 0.02% to keep the misclassification rate below 10% (Fig. 2 C). For more sensitive cluster/clade/grade-level identification, we recommend a misclassification rate below 5%, which requires a minimum *Prochlorococcus* abundance of 0.15% and *Synechococcus* abundance of 0.03% of all classified reads (Fig. 2 A and C).

**Figure 2.**
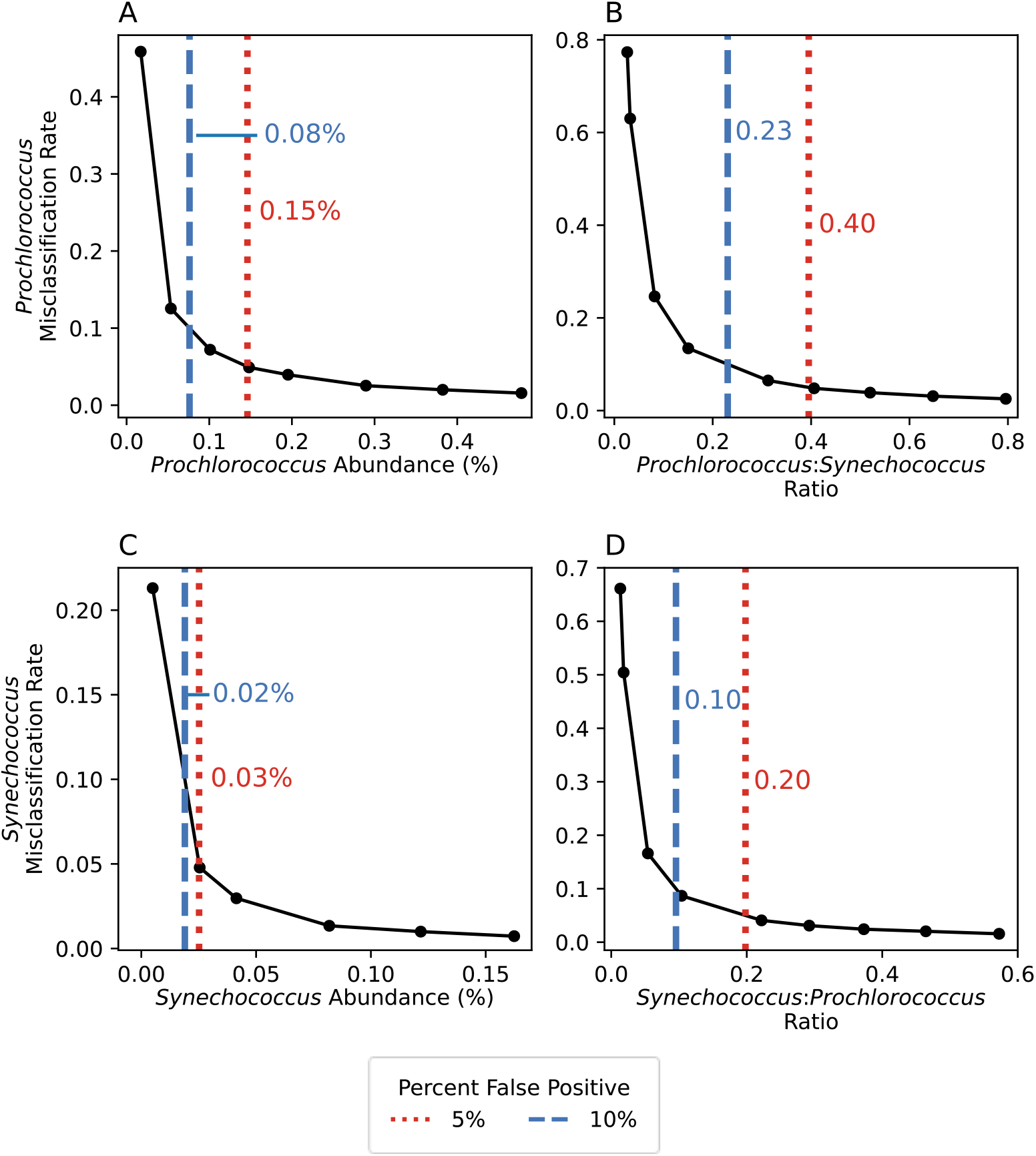
Misclassification rates and limit of detection for *Prochlorococcus* and *Synechococcus*. Misclassification rates examined across (A) *Prochlorococcus* abundances, (B) *Prochlorococcus:Synechococcus* ratios, (C) *Synechococcus* abundances, and (D) *Synechococcus:Prochlorococcus* ratios. Detection limits were determined at 5% (red dotted line) and 10% (blue dashed line) misclassification thresholds.

To establish the minimum *Prochlorococcus*:*Synechococcus* ratio and vice versa, we simulated samples with varying proportions of the two genera and processed them through the classification workflow. False positive rates were calculated by dividing misclassified reads by the total classified reads for each taxon. To keep the *Prochlorococcus* misclassification rate below 10%, the *Prochlorococcus*:*Synechococcus* ratio must be greater than 0.23 (Fig. 2 B). Similarly, to maintain a *Synechococcus* misclassification rate below 10%, the *Synechococcus*:*Prochlorococcus* must be greater than 0.10 (Fig. 2 D). For a more sensitive classification with a 5% misclassification rate, the required ratios are 0.40 for *Prochlorococcus*:*Synechococcus* ratio and 0.20 for *Synechococcus*:*Prochlorococcus* (Fig. 2 B and D).

### Cluster/Clade/Grade Accuracy

To evaluate the workflow’s accuracy in estimating the composition of different *Prochlorococcus* and *Synechococcus* clusters/clades/grades, we processed simulated samples with equal proportions of all clusters/clades/grades with and without marine heterotrophic bacteria, following the composition outlined in the Zenodo file (ProSynTax-workflow_benchmarking_composition.tsv)^37^. Specifically, we generated simulated samples containing only the respective picocyanobacterium, as well as mixtures where the respective picocyanobacterium was present at 2% with 98% marine heterotrophic bacteria, and at 1% with 98% marine heterotrophic bacteria plus 1% of the other picocyanobacterium.

The classification results for both *Prochlorococcus* and *Synechococcus* closely matched the expected values, except for a false positive classification of *Synechococcus* cluster 5.1B – V, which was absent from the simulated dataset, and a slight increase in false negative misclassification of *Prochlorococcus* grade LLVIII (Fig. 3 A) and *Synechococcus* cluster 5.1A – III (Fig. 3 B). The discrepancy of the latter is likely due to their underrepresentation in the genome dataset, which contains only five reference genomes for each cluster/grade.

**Figure 3.**
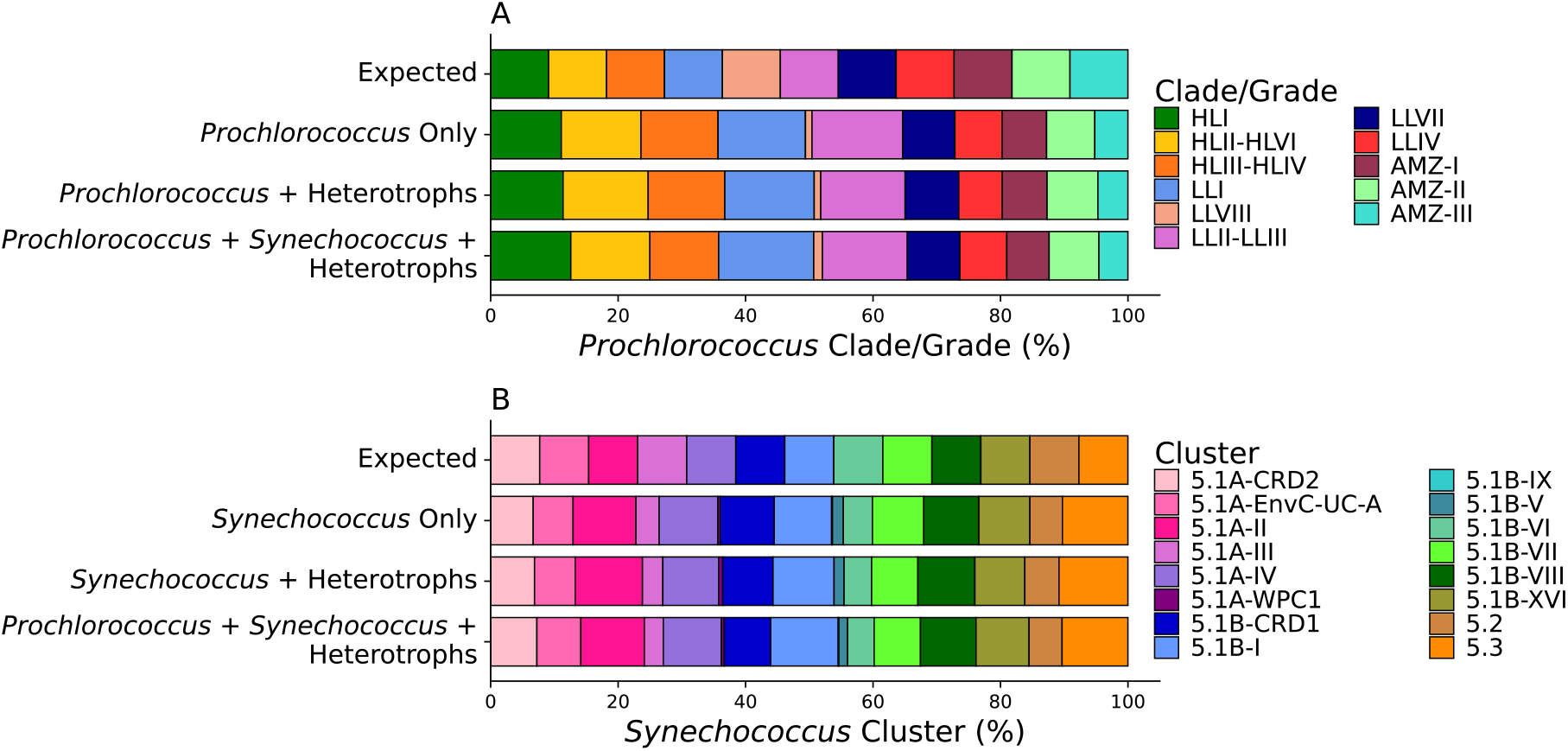
Accuracy of *Prochlorococcus* and *Synechococcus* cluster/clade/grade classification. Comparison of expected and classified percentages of *Prochlorococcus* (A) and *Synechococcus* (B) clusters/clades/grades in simulated samples of the respective picocyanobacterium alone, the respective picocyanobacterium at 2% with 98% marine heterotrophic bacteria, and the respective picocyanobacterium 1% with 98% marine heterotrophic bacteria and 1% of the other picocyanobacterium.

### Validation with field data

To evaluate the accuracy and effectiveness of ProSynTax and the associated workflow on field data, we analyzed the fraction of classified *Prochlorococcus* reads relative to the total reads (classified + unclassified) and assessed the clade/grade diversity throughout a depth profile at Station ALOHA using metagenomic data from cruises HOT224-238^46^. We applied the 5% detection limit threshold from Figure 2, enabling reliable estimation of total *Prochlorococcus* read abundance at each depth (Fig. 4 A). We then classified clade/grade diversity at each depth and found predominantly high-light (HL) *Prochlorococcus* clades at <75m and low-light (LL) clades/grades at >125m (Fig. 4 B). These results are consistent with many studies on *Prochlorococcus* diversity at Station ALOHA^22,47–49^, demonstrating that the ProSynTax dataset and workflow accurately capture ecological patterns consistent with previous studies.

**Figure 4.**
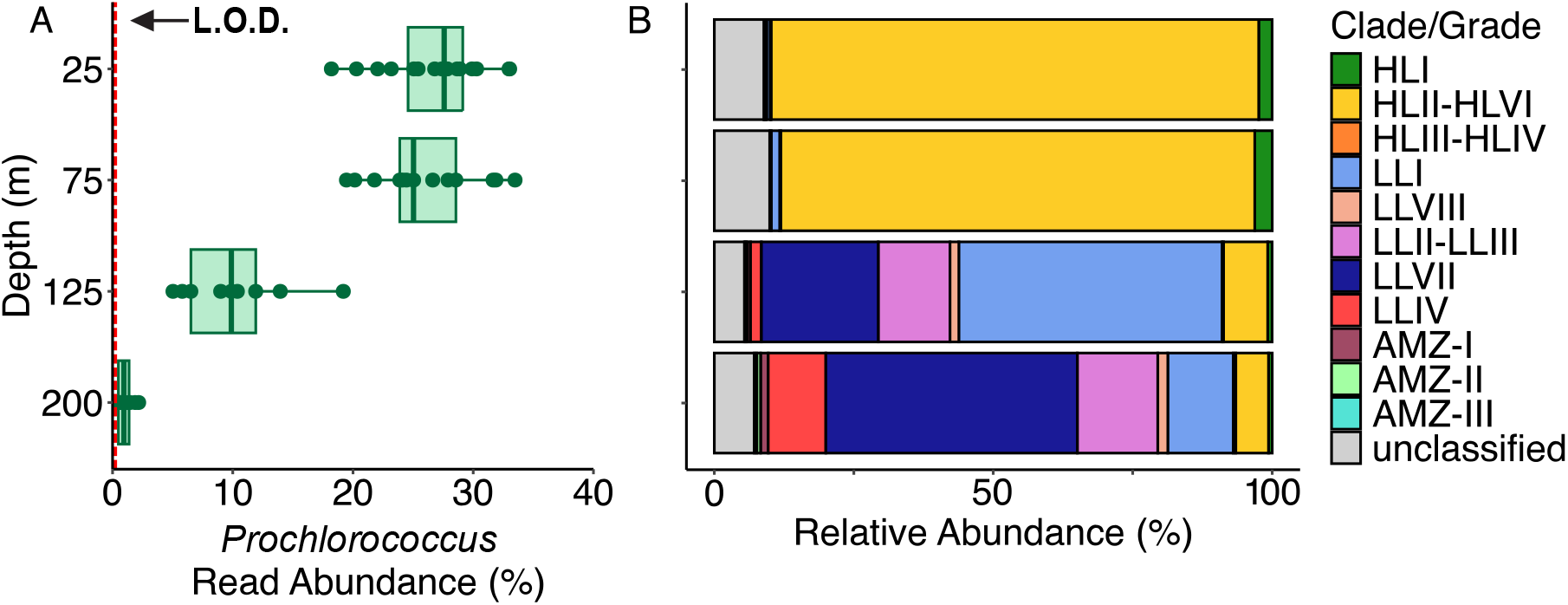
*Prochlorococcus* clade/grade diversity at STATION ALOHA. Metagenomes from research cruises HOT224-238^46^ were used to calculate (A) the fraction of classified *Prochlorococcus* reads relative to the total reads (classified + unclassified), including only samples above the 5% detection limit threshold (L.O.D., red dotted line, Fig. 2) and (B) the fraction of each classified clade/grade reads relative to total *Prochlorococcus* reads, calculated using estimated genome equivalents at each depth.

## USAGE NOTES

The ProSynTax dataset and workflow assigns taxonomic classifications to metagenomic reads based on the taxonomic hierarchy in ProSynTax and uses these classified reads to estimate the genome equivalents of *Prochlorococcus* and *Synechococcus* clusters, clades, or grades within a metagenome sample. This dataset can be customized by allowing users to add new genomes as needed as well as modify the taxonomy in order to improve classification accuracy and adapt the workflow to specific research needs.

To setup and run the ProSynTax workflow, first clone the ProSynTax workflow repository into a suitable directory. Then, download the protein dataset index file (ProSynTax_v1.1.fmi), the names (ProSynTax_names.dmp), the nodes (ProSynTax_nodes.dmp), and the diamond blast database (CyCOG6.dmnd) from the Zenodo repository^37^. Once the repository is cloned and the dataset files are in place, update the input files to specify the correct file paths and directories. To execute the workflow, navigate to the scripts directory and run the run_classify_smk.sbatch script.

The workflow begins by trimming low-quality regions and removing adapter sequences from paired-end FASTQ files using BBDuk v39.18^27^, followed by read classification with Kaiju v1.10.1^26^. A custom Python script^37^ summarizes raw read counts by genus, after which reads identified as *Prochlorococcus* and *Synechococcus* are extracted into separate FASTA files using Seqtk vr82^50^. To correct for genome length variation across clusters/clades/grades, raw reads are aligned to single-copy core genes (SCCG) from the CyCOG v6.0 database^21^ using DIAMOND Blastx v2.1.11^51^. Genome equivalents are estimated by normalizing total SCCG read residues to the average SCCG length.

It is recommended to filter out samples with low *Prochlorococcus* or *Synechococcu*s abundance and to adjust parameters when working with low-coverage samples. Samples showing high proportions of unclassified reads or significant taxonomic imbalances may also benefit from parameter tuning. For broader *Prochlorococcus* ecotype comparisons, a 10% false positive threshold is generally sufficient, whereas a stricter 5% threshold is recommended for cluster/clade/grade delineations (Fig. 2). For detailed guidance, comprehensive tutorials are available on GitHub (https://github.com/jamesm224/ProSynTax-workflow).

## CODE AVAILABILITY

All ProSynTax files are available on Zenodo^37^ Code and documentation for ProSynTax associated workflow, along with the accompanying technical validation are available on the GitHub repository: https://github.com/jamesm224/ProSynTax-workflow

Code and documentation for Genome Closing workflow are available on the GitHub repository: https://github.com/konnorve/HIFI-genome-closing-improved.git

## AUTHOR CONTRIBUTIONS

PMB, MAD, AC, SWC, JIM, NNV conceived and designed experiments. PMB and JWB isolated cultures and AC and PMB maintained cultures. AC, KGC, SMP, ES, CB, and AAA grew cultures and performed DNA extractions. NNV, PMB, KVM, and AC closed genomes and submitted data to repositories. JIM, PMB, and NNV developed code, workflow, and dataset. NNV and JIM performed validation of the dataset and MAD, AC, and PMB analyzed validation results. All authors contributed to writing and editing the manuscript.

## ACKNOWLEDGEMENTS

We thank the crews and research teams of the KM0915, MV1015, KM1217, KM1309, and KOK1404 cruises on the research vessels R/V Kilo Moana, R/V Melville, and R/V Káimikai-O-Kanaloa for their support in facilitating the *Prochlorococcus* isolations used in this study. This work was supported by grants from the National Science Foundation (OCE-1153588 to S.W.C.; DBI-0424599 to S.W.C.; OCE-2048470 to P.M.B.), the Gordon and Betty Moore Foundation (GBMF495 to S.W.C.; GBMF4511 to S.W.C.), the Robert and Ardis James Foundation to S.W.C., and the Simons Foundation (Life Sciences Project Award ID 337262, S.W.C.; SCOPE Award ID 329108, S.W.C.; Marine Microbial Ecology Postdoctoral Fellowship 984601, M.A.D.). This paper is a contribution from the Simons Collaboration on Ocean Processes and Ecology (SCOPE).

